# The functional foundations of episodic memory remain stable throughout the lifespan

**DOI:** 10.1101/2020.01.30.926469

**Authors:** D. Vidal-Piñeiro, MH. Sneve, IK. Amlien, H. Grydeland, AM. Mowinckel, J.M. Roe, Ø. Sørensen, LH. Nyberg, KB. Walhovd, AM. Fjell

## Abstract

It is suggested that the functional mechanisms behind specific forms of cognition, particularly episodic memory, may be dynamic over the lifespan and that cognitive preservation or decay in older age thus relies on age-specific mechanisms such as compensatory processes. Here instead, we tested whether the functional foundations of successful episodic memory encoding adhere to a principle of lifespan continuity, shaped by developmental, structural and evolutionary influences. We identified the generic lifespan patterns of memory encoding function across the brain (n = 540; age range = 6 – 82 years). The lifespan trajectories of brain activity were organized in a topologically meaningful manner and aligned to fundamental aspects of brain organization, such as large-scale connectivity hierarchies and evolutionary cortical expansion gradients. None of the normative trajectories of encoding function was solely determined by late-life patterns of activity, but rather showed continuities across development and adulthood. Inter-individual differences in activity in age-sensitive regions were predicted by general cognitive abilities and variation in grey matter structure, which are core variables of cognitive and structural change throughout the lifespan. Altogether, the results provide evidence for the lifelong continuity of the functional foundations of episodic memory which are bounded by both brain architecture and core mechanisms of cognitive and structural change over life. We provide novel support for a perspective on memory aging in which maintenance and decay of episodic memory in older age needs to be understood from a comprehensive life-long perspective rather than as a late-life phenomenon only.

**Significance statement:** It is suggested that cognitive function in older age largely relies on late-life specific mechanisms such as compensatory processes. In contrast, here we tested whether and to what degree brain activity during episodic memory encoding adheres to fundamental principles of life-long brain organization and continuity. The results revealed that generic lifespan trajectories of memory encoding function were not specific to late-life. Instead, the age-trajectories showed a continuity through life, were related to fundamental features of brain structure and cognition and to functional and evolutionary hierarchies. We argue that rather than focusing on older-age specific mechanisms, a framework that takes lifespan mechanisms of cognition and brain anatomy into account is necessary to understand episodic memory vulnerability in older age.

## Introduction

Over the last few years, research has demonstrated that the structural foundations of general cognitive abilities are largely constant throughout life (1, 2), being embedded into fundamental aspects of brain organization as captured by evolutionary expansion patterns or connectivity gradients (3, 4). However, it is unknown whether the functional foundations supporting specific forms of cognition are equally stable or rather dynamic through life. In supporting signature aspects of human cognition like autonoetic consciousness and future thinking (5, 6), episodic memory represents a crucial ability in everyday function. Its vulnerability, particularly in old age, has thus attracted much research effort, leading researchers to postulate distinct age-specific mechanisms to explain brain-behavior correlates at different periods in life (7–9). Yet, lifespan researchers have emphasized integrative accounts of lifelong changes in cognitive abilities - and episodic memory in particular –, in which development and decay of brain structure often represent a key fundament for brain function and cognitive change (10–14). Using a novel and multifaceted analytic approach, we tested the continuity vs. age-specificity in the functional mechanisms supporting episodic memory and how these are related to fundamental variations in brain structure and cognition throughout the lifespan. We aimed to assess whether episodic memory function represents fundamental aspects of life-long brain organization and continuity, similar to what has been established for general cognitive abilities.

We delineated the lifespan trajectories associated with episodic encoding success, using fMRI data from 540 healthy individuals from 6 to 82 years during an incidental source-item encoding task (**Fig. 1a-c**). Region-wise encoding activity throughout the entire brain was fit to age using generalized additive models (GAM) and the most characteristic lifespan trajectories were established based on a clustering approach (**Fig. 1d-g**). Encoding was indexed using a source vs. item memory contrast that isolates the binding aspects of episodic memory and is highly sensitive to age. We reasoned that support for an account of lifelong continuity in the brain foundations of episodic encoding would require that: a) The lifespan trajectories of encoding activity cluster into known networks with a meaningful function and topology; b) The clusters of activity trajectories are embedded in fundamental aspects of brain organization, namely evolutionary-related cortical expansion and functional connectivity gradients (3, 15, 16). c) The trajectories are not determined by age-specific profiles of activity at middle or older age, i.e. defined by increased/decreased activity in late-life only. Rather, lifespan trajectories of encoding function are predominantly determined by developmental profiles. d) Interindividual variations of activity in developmentally sensitive clusters are related to core cognitive abilities and structural features that govern brain and cognitive variability in childhood and older age.

**Fig. 1.**
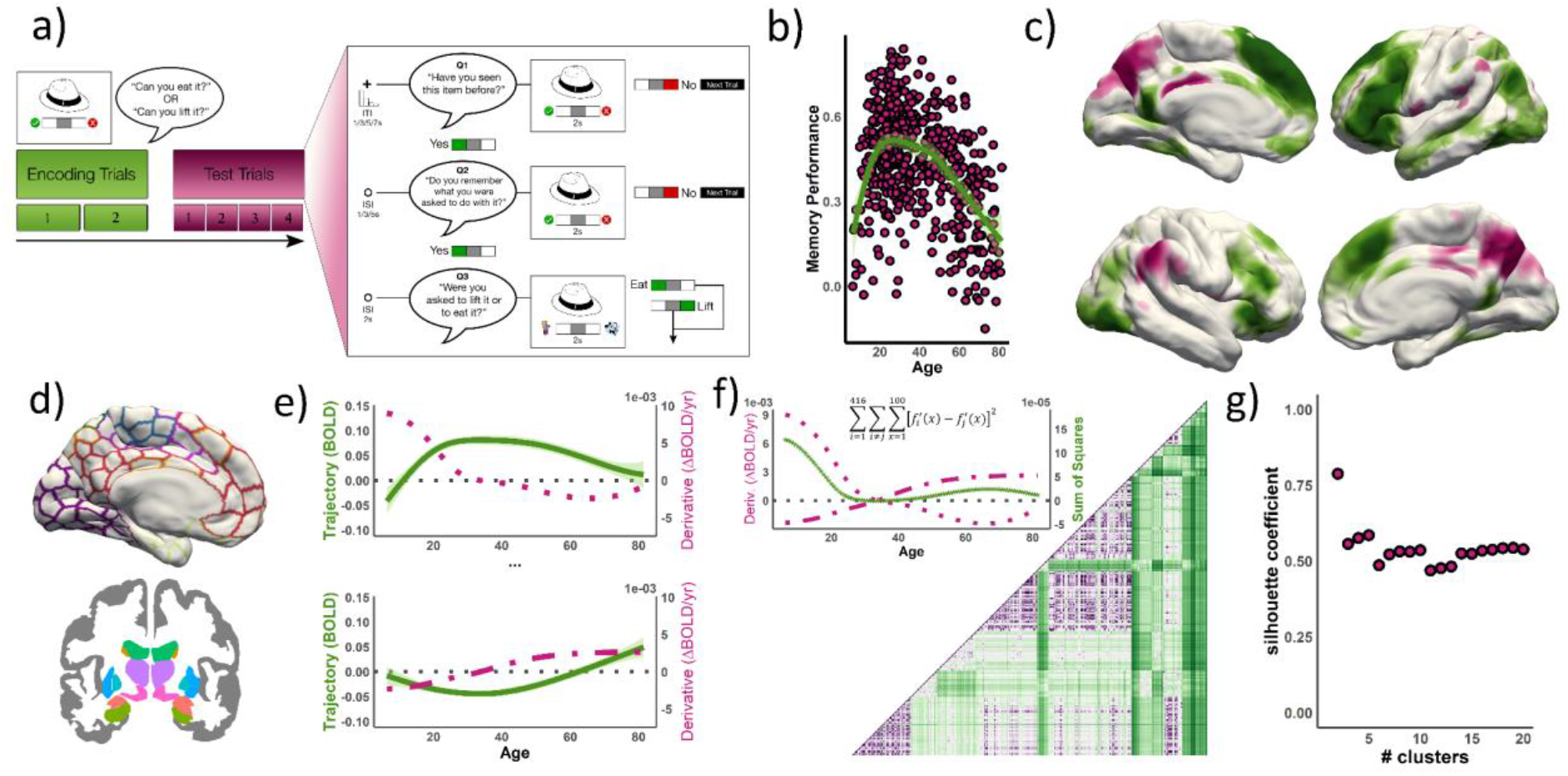
Experimental scheme. a) Experimental design; adapted from (17). See **SI Methods** for more information. b) Source memory performance across the lifespan sample (proportional, corrected for guessing) c) Subsequent memory contrast (BOLD_S>i_: source vs item memory encoding) for the entire sample (n = 540). **See SI Methods and Results.** Green and purple regions denote positive and negative subsequent memory effects, respectively. d-g) Lifespan Parcellation pipeline. Brain clusters characterized by different ‘canonical’ trajectories were established from subsequent source memory encoding effects across life. d) 416 ROIs were selected corresponding to the “Schaefer” and “fsaseg” cortical and subcortical atlases. e) For each ROI, we fitted a lifespan trajectory to encoding activity and computed its derivative. f) To group regions with similar age trajectories of encoding function, we obtained a dissimilarity matrix by computing the distance between each pair of ROIs’ derivatives - based on a least square sum. Regions without evidence of subsequent memory activity (n = 74) were removed from the dissimilarity matrix. g) Regions showing subsequent memory activity were fed into a k-medoids algorithm and the optimal partition (k = 5) was established by the silhouette width coefficients.

## Results

### Delineation of lifespan trajectories of encoding activity

As expected, age was related to memory performance (F = 37.4, p < .001, edf [estimated degrees of freedom; index of curve complexity] = 4.5; **Fig. 1c**), revealing an inversed U-shape lifespan trajectory. See complete behavioral results in **Fig. S1** and **Table S1**.

We reduced the number of regions in a data-driven manner by clustering the brain based on the pairwise similarity of the lifespan trajectories (derivatives) of episodic encoding function. We used a k-means clustering algorithm, which yielded an optimal partition at k = 5 with an average silhouette coefficient of .59. **Fig. 2** displays the resulting 5-partition arrangement of cortical and subcortical regions based on the canonical trajectories of encoding activity during the lifespan. For each cluster, a canonical trajectory refers to the mean lifespan trajectory against which the other trajectories are compared and adhere to. See in **Fig. 3** the effects of age, edf, and mean activity per ROI grouped by cluster. Note that by using a distance matrix based on the derivatives, we partitioned the brain solely from the lifespan trajectories, disregarding the intercept (i.e. mean activity). Thus, regions are grouped together if their episodic encoding activity shows the same age-relationships (e.g. two regions with similar lifelong trajectories will group together regardless of whether they show positive or negative memory effects).

**Fig. 2.**
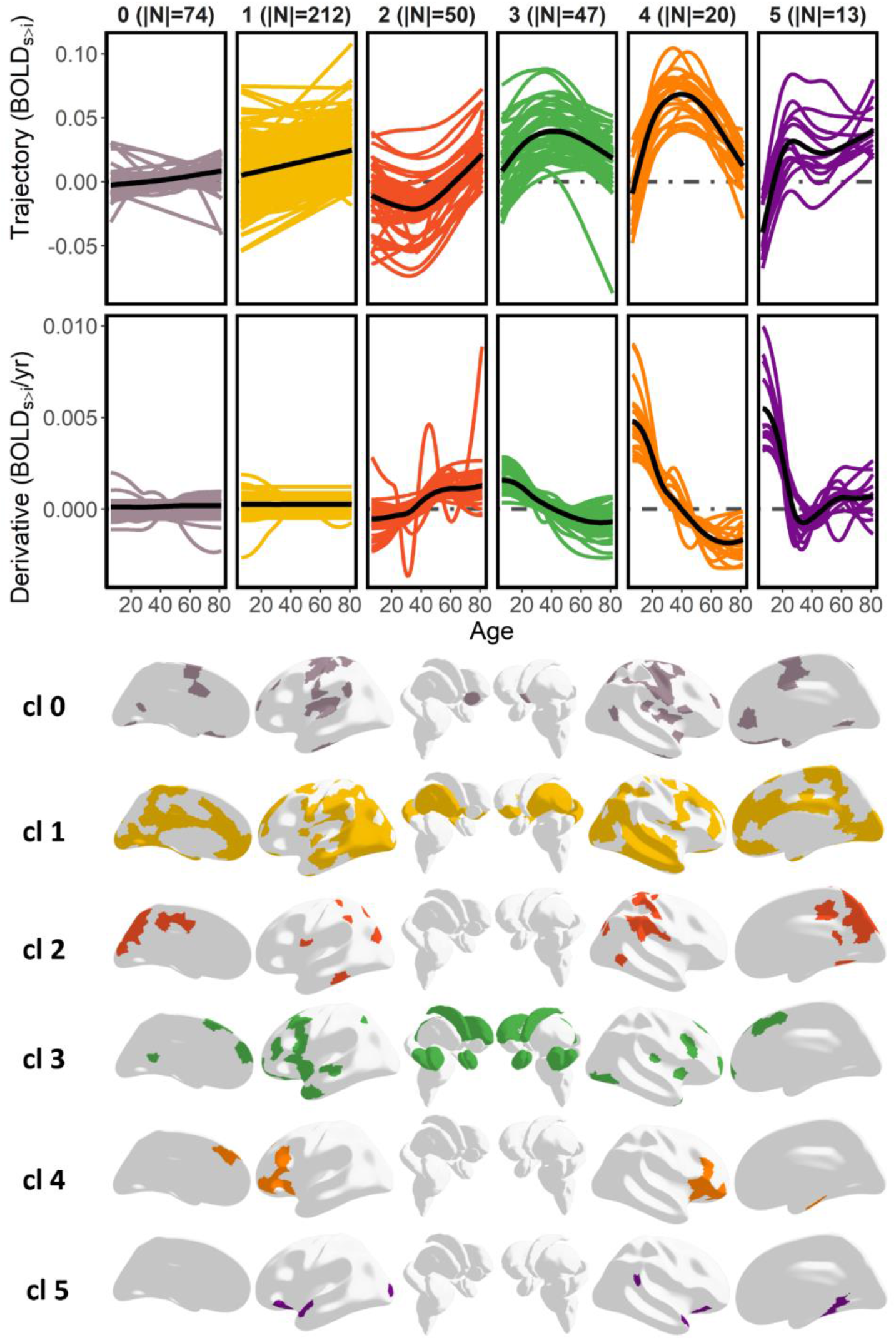
Cluster solution based on the derivatives of the lifespan trajectories of encoding activity. Upper panel: Lifespan trajectories and the derivatives of encoding activity grouped by cluster. Lower panel: ROI assignment by cluster. BOLD_S>i_ = Subsequent source vs. Item memory fMRI contrast. Note that cluster 0 was defined prior to the clustering analysis as regions not showing subsequent memory effects.

**Fig. 3.**
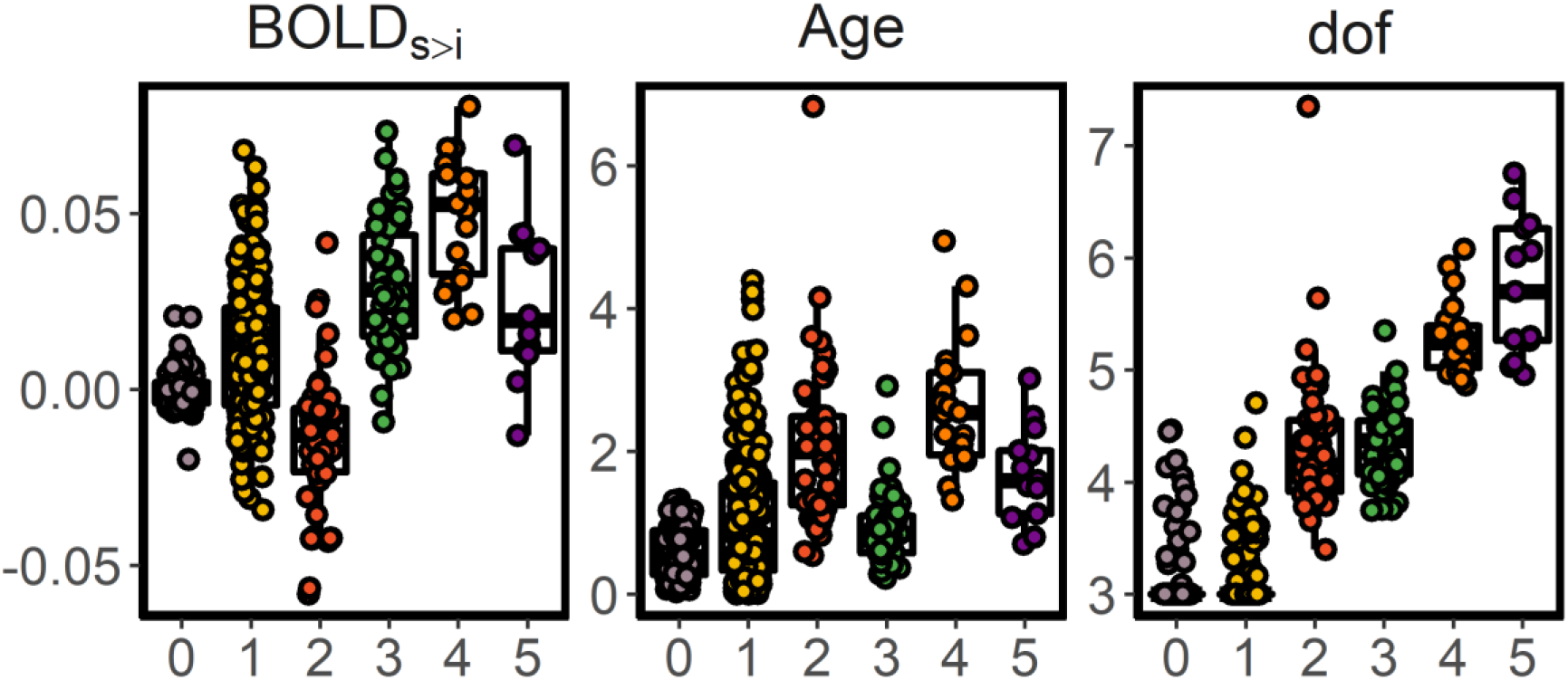
Mean BOLD activity for the source-v-item memory contrast (BOLD_S>I_), age effects (−log10(p)), and estimated degrees of freedom (range = 3 - 9) for each ROI grouped by cluster.

An initial group of regions (cluster 0) was obtained prior to the partition algorithm (|N| = 74), which included regions that did not show evidence of activity associated with encoding success throughout the lifespan. Most other regions were assigned to cluster 1 (|N| = 212), which encompassed large parts of cortex and subcortex including the posterior hippocampi. The trajectories of cluster 1 tended to exhibit weak monotonic increments of activity throughout the lifespan. Cluster 2 consisted of |N| = 50 regions, which were located almost entirely in the posteromedial and in the inferior parietal lateral cortices. Cluster 2 showed a canonical U-shape lifespan trajectory and consisted of regions that showed negative memory effects during young adulthood. Cluster 3 consisted of |N| = 47 regions from bilateral prefrontal and subcortical regions, including the anterior hippocampi, that mapped onto a weak inverted-U shape trajectory. Most of the regions exhibited positive subsequent memory effects during young and middle adulthood. Most of the |N| = 20 regions of cluster 4 mapped to the inferior frontal gyrus, bilaterally, extending also to the left superior frontal cortex and the right parahippocampal gyrus. Episodic encoding activity in cluster 4 showed a steep inverted U-shaped trajectory over the lifespan. All these regions exhibited positive subsequent memory effects during young and middle adulthood and were significantly related to age (all ROIs p < .05). Finally, cluster 5 consisted of |N| = 13 regions and exhibited a canonical *developmental* trajectory with activity increasing in childhood before reaching a plateau that lasted throughout adulthood. Cluster 5 included anterior temporal, pars orbitalis regions bilaterally as well as parts of the right temporoparietal and parahippocampal cortices. Overall, the results showed a continuity of the canonical trajectories throughout the lifespan as a) patterns of activity developed and decayed at younger and older age, respectively (clusters 2-4); b) developed at younger age and later stabilized (cluster 5) or, c) showed a monotonical pattern through the entire life (cluster 1). Critically, none of the trajectories exhibited late-life profiles of activity with distinct patterns emerging at middle or older age.

### Relationship of lifespan encoding clusters with cognitive function

We next tested whether variations of activity in the encoding clusters related to interindividual differences in core cognitive functions, as indexed by Matrices Reasoning and Vocabulary scores (18). Further, we tested the relationship between cluster activity and memory performance as indexed both by task performance in the fMRI task as well as by an external verbal recall task (CVLT learning) (19). We ran parallel GAM models with age and the cognitive tests as smoothing terms, principal component analysis (PCA)-based cluster activity as outcome, and sex as a covariate. See **Fig. 4** for a visual representation of the relationship between BOLD activity and cognitive function. Activity in cluster 4 was significantly associated with better performance on the fMRI task (F = 7.8, pFDR [False Discovery Rate corrected using the Benjamini–Yekutieli procedure (20); n = 24] = .006, edf = 2.2) and matrix scores (F = 10.0, pFDR = .04, edf = 1.1) while CVLT learning scores were close to significance (F= 4.1, pFDR = .05, edf = 2.2). Activity in clusters 5 and 3 was associated with better vocabulary and matrix scores), respectively (F = 6.6, pFDR = .01, edf = 2.7; F = 11.4, pFDR = .03, edf = 1.3. The remaining comparisons did not pass the significance threshold. Note that the relationship with vocabulary scores in cluster 5 flattens with higher cognitive performance suggesting that the observed non-linear association is mostly driven by the younger participants. See complete stats in **Table S2**. The results indicate that activity in developmentally sensitive clusters is linked to performance in established core functions known to drive cognitive change throughout the lifespan.

**Fig. 4.**
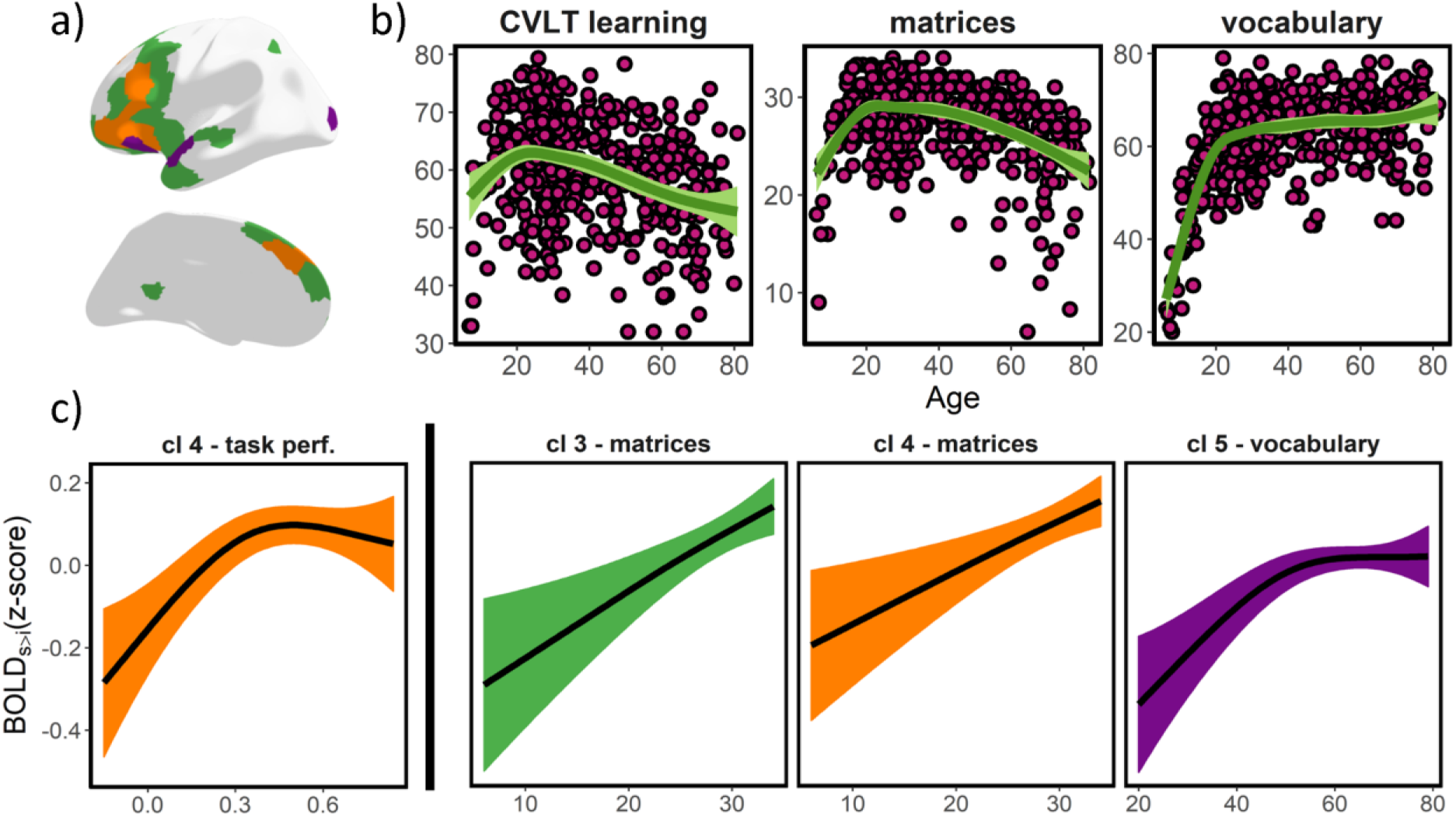
Episodic encoding activity – cognitive function relationships. a) Topography for cluster 3, 4, 5 (green, orange, purple; left hemisphere only). b) Lifespan trajectories of CVLT learning, Matrices, and Vocabulary scores. c) Significant (FDR-corrected) relationships between cluster activity and cognitive function, controlling for age and sex. x-axis represents test scores and y-axis represents (source-v-item) encoding activity. Ribbons represent 95% confidence intervals. Note that activity (y-axis) values are derived from a PCA (within cluster), and thus demeaned. For clusters 3-5, higher values represent stronger positive memory effects (see **Fig. 2**). BOLD_S>i_ = Subsequent source vs. item memory fMRI contrast.

### Relationship of lifespan encoding clusters with large-scale modes of GM variation

Next, we assessed whether interindividual differences in activity in developmentally sensitive clusters were associated with core features of structural brain variability throughout the lifespan. We used a linked Independent Component Analysis (ICA) (21, 22) to derive modes of grey matter (GM) variation using three different modalities: cortical thickness and area based on cortical surface reconstructions (23), and volume from a voxel-based morphometry (VBM) protocol (24). As described in (25), we identified two GM components that showed a strong relationship with age. The first, IC_GM_1, represented a dominant whole-brain mode of variation, explaining ~25% of the structural variance across individuals. This component showed a monotonic decrease in GM across the lifespan. Age explained ~86% of the IC_GM_1 variance as assessed post-hoc with GAM. The second, IC_GM_2, explained ~5% of the total GM variance and loaded heavily on prefrontal and parietal heteromodal areas. IC_GM_2 exhibited an inverse U-shape trajectory with age, which explained 46% of the component’s variance. See additional details in **SI methods, SI results** and, **Fig. S2-3**.

GAM analysis – using age and GM variation as smoothing terms and sex as covariate - revealed that interindividual differences in GM captured by IC_GM_2 related to higher encoding activity in clusters 4 and 5 (F= 11.3, pFDR = .01, edf = 1; F = 23.9, pFDR < .001, edf = 1, respectively). IC_GM_2 network mapped onto areas susceptible to normal and abnormal developmental and aging changes (25). In addition, encoding activity in cluster 5 was associated with GM loadings in IC_GM_1 (F = 5.7, pFDR = .01, edf = 2.5) (**Fig. 5**). See full stats in **Table S3**. Note that during development, the relationship between GM indices such as cortical thinning and cognition is typically negative (26), which can explain the negative relationship between IC_GM_1 variation and activity, which exists only for high GM loads. Thus, the results suggest that cluster activity is constrained and supported by the development and decay of large modes of GM variation throughout the lifespan.

**Fig. 5.**
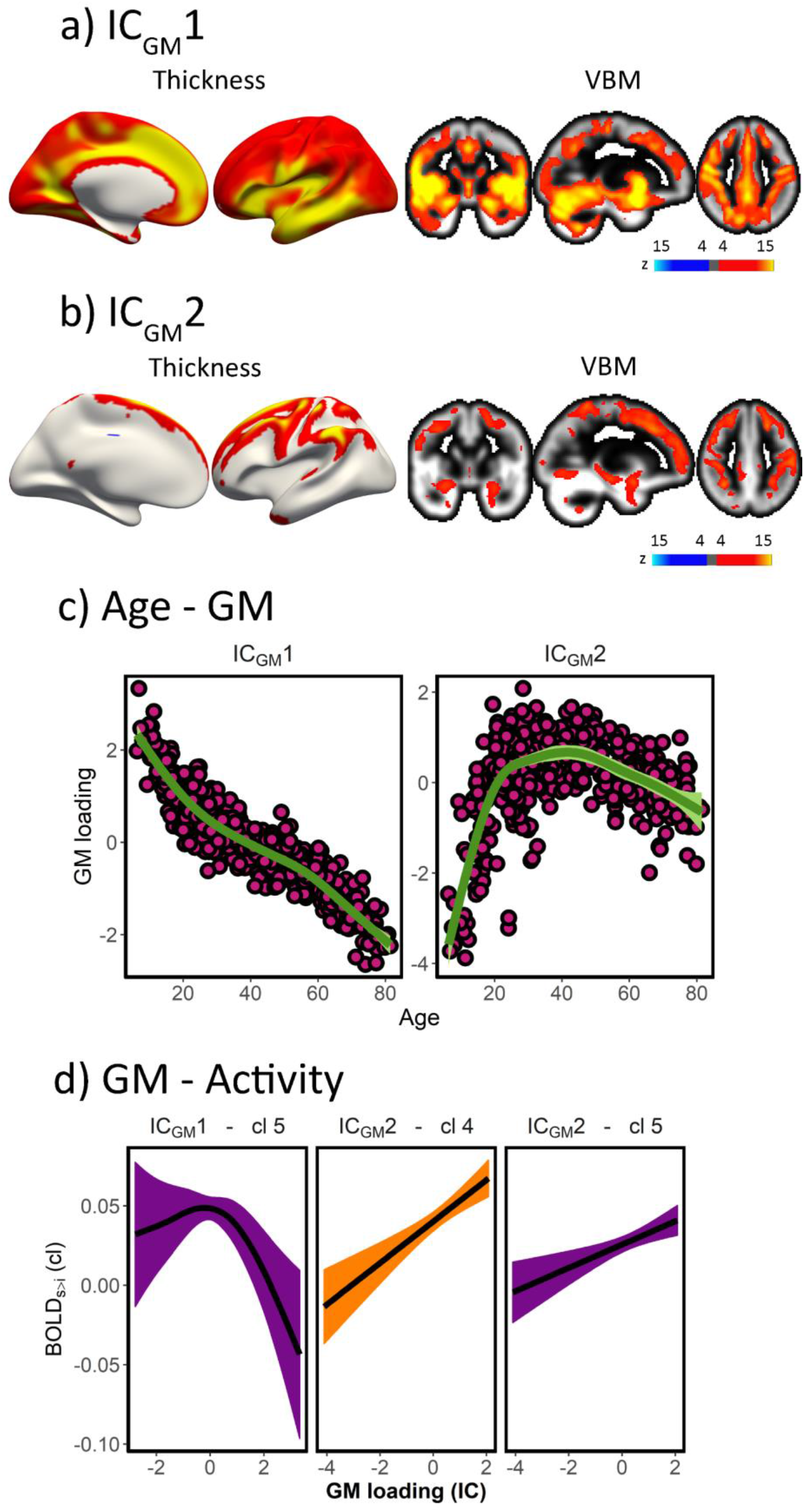
Activity – GM variation. a, b) Modes of GM variation with practical (r^2^ > .15) age significance. Area loadings not displayed; see **Fig. S3**. c) Relationship between age and modes of GM variation. d) Relationship between GM variation and cluster activity (sex, age-corrected). Only significant relationships after FDR-correction are shown. x-axis represents GM loadings and y-axis represents activity. Ribbons represent 95% confidence intervals. Note that activity (y-axis) values are derived from a PCA and thus centered to 0. For clusters 3-5, higher values represent stronger positive memory effects. See **Fig.2**.

### Topological relationship of lifespan partitions with functional and evolutionary hierarchies

Finally, we tested whether the lifespan trajectories of encoding activity were embedded in fundamental aspects of brain organization as indexed by flexibility, the principal gradient of functional connectivity, and cortical expansion through evolution. Flexibility indexes the degree to which a region participates in multiple cognitive components, likely by binding and integrating specialized brain networks (16). The principal gradient of functional connectivity represents an overarching organization of large-scale connectivity that reflects a functional hierarchy from perception/action (in sensorimotor areas) to abstract cognitive functions (in the default mode network) (3). The expansion index reflects the degree to which a region has grown in size between macaque and humans (15, 27). These three measures reflect different fundamental aspects of brain organization in which higher values reflect diminished constraints of sensory and structural input and increased capacity to support a wider array of different tasks such as higher-order cognition (4, 28, 29). **Fig. 6** presents the topological relationship between clusters - based on lifespan trajectories of encoding function – and the functional and evolutionary hierarchical maps. Results showed that cluster 4 encompassed regions characterized by high flexibility (pFDR = .002), high macaque to human expansion (pFDR = .04), and aligned at the apex of the functional connectivity hierarchy (pFDR < .001) while cluster 3 was characterized by regions aligned at the apex of the functional connectivity hierarchy (pFDR < .001). See complete stats in **Table S4**.

**Fig. 6.**
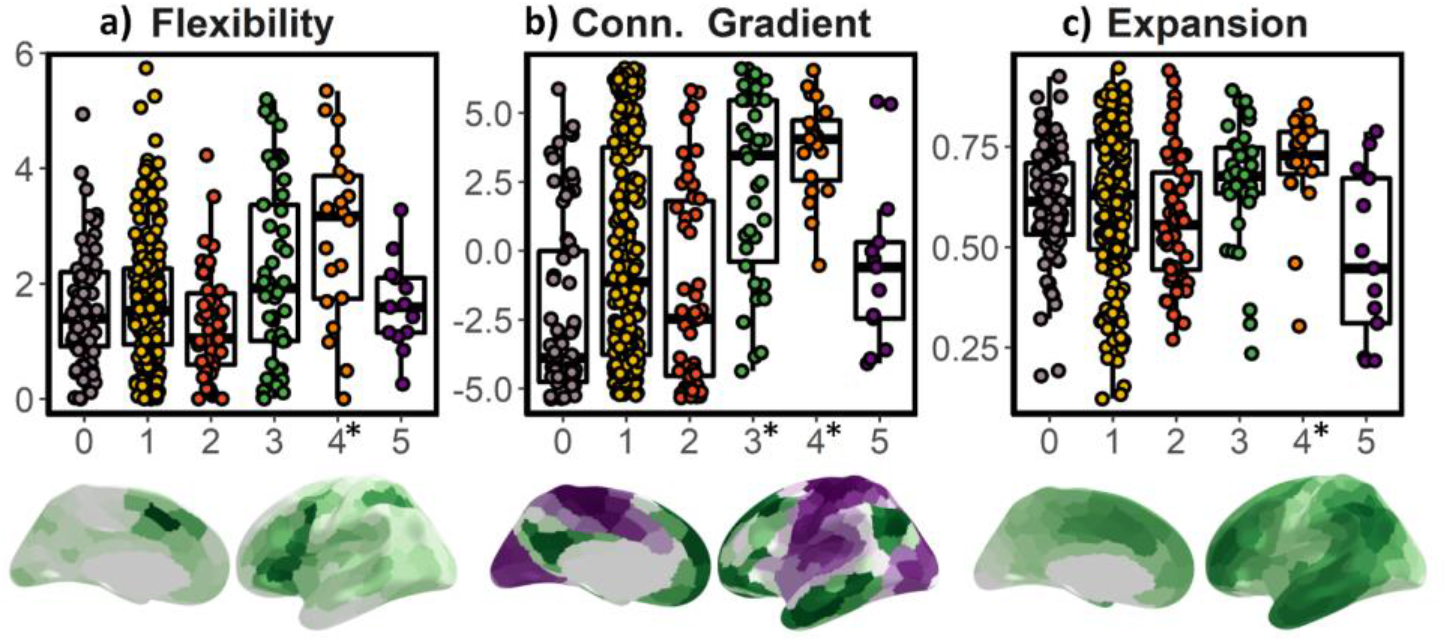
Topological relationship between encoding clusters and functional and evolutionary hierarchy. a) Flexibility (16), b) specialization determined by the principal gradient of functional connectivity (3), and c) macaque-human expansion (15, 27). Y-label represents the number of recruited components, distance (mm), and normalized (0-1) expansion, respectively. Conn. Gradient = principal gradient of functional connectivity. * denotes significance after permutation testing.

## Discussion

Overall, the results suggest that episodic encoding activity exhibits a continuity from childhood to older age, supported by age-sensitive features of brain structure and general cognitive functions. Existing evidence suggests that differences in late-life general cognitive function and brain structure are shaped by early-life influences (1, 2). Here we show that this principle also applies to the brain activity underlying episodic encoding success and that episodic memory trajectories are also determined by fundamental aspects of brain organization such as cognitive flexibility, functional connectivity, and evolutionary cortical expansion. Critically, we did not find evidence of age-specific profiles of activity emerging at middle or older age. Rather, memory encoding trajectories were always influenced by developmental profiles. While some cortical regions showed higher encoding-related activity in older age, they invariably corresponded to monotone increments originating in childhood, suggesting that these patterns did not reflect compensatory responses or dedifferentiation processes appearing at old age. The findings refer to general patterns of activity and thus do not necessarily preclude the existence of specific mechanisms supporting successful encoding in older age, but limit its extent to small subsamples of participants or to processes with unspecific spatial distribution. Thus, the results suggest that episodic memory encoding is related to fundamental brain characteristics in much the same way as general cognitive function, and that successful episodic memory encoding in higher age should be understood in a lifespan perspective. The specific results are discussed below.

### Identification of canonical lifespan trajectories of episodic encoding activity

In the present study, the brain was parcellated based solely on the shape of the lifespan trajectories of encoding activity. This novel, data-driven approach revealed several clusters characterized by unique lifespan trajectories that mapped to well-characterized functional and evolutionary patterns in the brain. This observation is illustrated by the inverted U-shaped trajectory of cluster 4 that mostly included bilateral inferior and superior prefrontal regions. Prefrontal cortex activity is thought to reflect a set of cognitive control operations that support the encoding of discrete memory traces (30). The inverse U-shape trajectory of encoding success through life was closely aligned with those of fluid cognitive abilities and prefrontal brain structure integrity (31, 32) and fits well with the proposition that strategic components of episodic memory undergo a protracted maturation in childhood and a pronounced decline in old adulthood (13). Indeed, we found that cluster 4 activity corresponded to higher fluid intelligence (matrix reasoning performance) and higher GM loadings in a frontoparietal heteromodal network. Cluster 4 regions map to a network characterized by protracted development, i.e. prefrontal cortex, and is characterized by marked changes in neurobiology including myelination, synaptic pruning and dendritic remodeling (33). This prefrontal activity captured by cluster 4 may, in part, reflect the maturation and decline of cognitive control components of encoding function, thus representing an important mechanism for both general and domain-specific cognitive change throughout life (10).

The cluster 5 trajectory was characterized by increasing episodic encoding activity during development and a plateau through adulthood. This pattern can only be revealed via a lifespan approach, as age-relationships were limited to childhood/adolescence, and the purely developmental nature of the cluster would have been concealed without mapping across a wider age-rage. This cluster included parts of the anterior lateral temporal cortex, the temporoparietal junction, and anterior inferior frontal regions. One can speculate that cluster 5 reflects a group of regions that are involved in high-level conceptual processes. The trajectories of this cluster mimic the lifespan trajectories for representational knowledge and map well to a network involved in semantic/conceptual processing (34–36). Cluster 5 activity may thus be particularly sensitive to the maturation of conceptual processing mechanisms. Higher cluster 5 activity further related to higher vocabulary scores and lower GM loadings of the first component. This GM component reflects, to a large extent, cortical thickness, and has a strong negative correlation with age, due to the steep rate of apparent cortical thinning during development (37). The maintenance of cluster 5 activity combined with the inverse U-shape trajectory of cluster 4 in prefrontal structures is compatible with the default-executive hypothesis of aging, which posits that with increasing age cognitive processes rely more strongly on semanticized mechanisms (38). Altogether the results also conform to the brain maintenance view to the extent that preserved cognition in aging relates to maintaining youthful brain structure, function, and neurochemistry (11) (e.g. **Fig. 5D**).

Lifespan variations of episodic encoding activity are linked to variations in GM integrity – likely capturing changes in myelin and dendritic arbors (39–41) – and to major mechanisms of cognitive change through the lifespan, namely fluid and crystallized abilities (10). Further, regions showing the most marked changes in episodic encoding activity through life are located in parts of the cortex characterized by strong expansion in primate evolution with a function less constrained by brain structure and sensory input, and hence, able to support a wider array of different task configurations (3, 16, 29). Previous work has shown that inter-individual differences in cortical morphometry in hotspot regions of expansion are related to general cognitive function (42), brain development (27), aging and Alzheimer’s Disease (42), and brain activity both during rest and task execution (4). The alignment with the different cortical organization maps suggests that the encoding activity in these regions represents cognitive elements that are continuously developing throughout life as well as being either uniquely human or at least disproportionally developed in humans. In return, these features might also confer a region with heightened vulnerability to the effects of age and disease (42, 43).

### Limitations

Two technical issues to consider relate to the clustering pipeline and the effects of motion on the results (e.g. cluster solution). The *k-*medoids algorithm is a data-driven clustering method. Different parcellations or dissimilarity matrices may yield different clustering solutions. Also, *k-*medoids is blind to any predefined lifespan trajectory and forces every ROI into a cluster. As such some ROIs might not fit well within a given cluster, yet it is unadvisable to exclude them based on this basis as it assesses similarity to the cluster, not to a predefined trajectory. Certainly, motion is strongly associated with age and affects BOLD signal although the impact is minor when using fMRI contrasts. Yet, motion correction is a double-edged sword as removing participants with high motion introduces sample bias while covarying motion out is also problematic as it relates to maturation and decline of brain structure (44).

Although we believe that it is very unlikely that in a given region and within our constrained experimental setup, different mechanisms of successful encoding are at play at different ages, a potential caveat arises by regional activity reflecting different supporting mechanisms of memory encoding during different periods in life. However, this issue remains largely hypothetical. With regard to the experimental setup, we predict the results will extend to other subsequent memory contrasts able to isolate binding mechanisms of episodic memory. Yet, it is unclear whether the results will generalize to situations characterized by intentional encoding and increased environmental support (45). Finally, the present study consists of cross-sectional, correlational data. Longitudinal studies are needed to characterize intraindividual trajectories of function, reveal lead-lag relationships, and uncover specific genetic and environmental influences on memory function trajectories. Ultimately, only longitudinal data will be capable of revealing the functional determinants of cognitive change throughout life (46).

## Conclusion

The study provides support in favor of stable functional foundations of episodic memory through life, from childhood to older age, instead of qualitatively different, age-specific, mechanisms. Variations in episodic memory were related to fundamental features of brain structure and cognition that characterized development and aging. We conclude that understanding memory vulnerability in older age requires a life-long comprehensive framework that considers normative cognitive, structural and functional aspects of memory function throughout the lifespan.

## Material and methods

### Participants

The final sample included 540 individuals (females = 366, age = 39.1 [SD = 18.5] years, age range = 6-82 years). The study was approved by the Regional Ethical Committee of South Norway. All participants ≥12 years gave written informed consent, all participants <12 years gave oral informed consent and, for all participants <18 years, written informed consent was obtained from their legal guardians. All participants were screened through health and neuropsychological interviews. See **SI Methods** for additional sample and exclusion criteria details. Participants’ data were discarded due to technical errors, faulty acquisitions or a low number of trials in a condition of interest (< 6 trials; n = 14).

### Experimental design and behavioral analysis

The experiment consisted of an incidental encoding task and a memory test after approximately 90 minutes. Both tasks took place inside the scanner. The experimental design is thoroughly described elsewhere (47, 48). For behavioral analysis, test trial responses to old items were classified as follows: source memory (Yes response to Q1 and Q2 and correct response to Q3); item memory (correct Yes response to Q1 and either a No response to Q2, or an incorrect response to Q3); or miss (incorrect No response to Q1). See **Fig. 1a** and **SI Methods** for further details.

### MRI acquisition and preprocessing

Imaging data were collected using a 20-channel head/neck coil on a 3T MRI (Skyra, Siemens Medical Solutions) at Rikshospitalet (Oslo). Each encoding run consisted of 134 volumes with the following functional imaging parameters: 43 transversally oriented slices were measured using a BOLD-sensitive T2*-weighted EPI sequence (TR = 2390 ms, TE = 30 ms, flip angle = 90⁰; voxel size = 3×3×3 mm; FOV = 224×224 mm; interleaved acquisition; GRAPPA factor = 2). Anatomical T1-weighted and gradient-echo field sequences were additionally acquired (see **SI Methods**).

The MRI dataset was converted to Brain Imaging Data Structure format (BIDS) (49) while cortical reconstruction and volumetric segmentation of the T1-weighted scans were performed with the FreeSurfer v.6.0 pipeline (http://surfer.nmr.mgh.harvard.edu/fswiki) (23). The fMRI analyses were constrained to a set of 416 ROIs covering the entire cortical and subcortical space. Subcortical ROIs regions were defined in native space based on the FreeSurfer automatic subcortical segmentation “*aseg”* with the additional division of the hippocampi along the anterior-posterior axis (50). Cerebellum was not included due to partial acquisition for this region in several participants. For the cortical surface, 200 ROIs were defined per hemisphere on each participants’ native reconstructed surface based on the Local-Global Parcellation of the Human Cerebral Cortex (51) (**Fig. 1d**).

fMRI data were processed using the “fMRIPrep” preprocessing pipeline (52). See **SI Methods** for a detailed description. The pipeline included skull-stripping, susceptibility distortions correction, motion correction, co-registration with the anatomical reference using boundary-based registration, and slice-timing correction. Post-*fMRIPrep* nuisance regression removed effects of estimated motion confounds (3 translations, 3 rotations, framewise displacement), and six *aCompCor* principal components derived from an eroded WM/CSF-mask. Data were high-pass filtered (128s cut-off) using a discrete cosine filter. Volume resampling was performed in a single interpolation step and was then sampled to each participants’ cortical surface space.

### fMRI analysis

First-level general linear models (GLM) were carried out with FSFAST. Briefly, percent signal change during source and item memory encoding was estimated and contrasted to produce estimates of episodic memory encoding, which were then averaged over voxels/vertices within ROIs. All between-subjects analyses were performed in R-environment (https://www.r-project.org/; v.3.5.2). When appropriate, significance values were corrected using false discovery rate (FDR) as implemented in (20) which controls for positive dependency amongst variables. We used *ggplot2*, *ggseg*, and *freesurfer* software for visualization (53, 54). See **SI Methods** for more details.

### Clustering of lifespan trajectories of encoding activity

In each ROI (n = 416), we fitted age on the episodic memory contrast using GAM models as implemented in the *vows* package. In each GAM, we fitted the activity values using age as the smoothing term (knots = 10) and sex as a covariate. The GAM models were re-run after excluding outlier values, defined as observations where residuals were >4 SD above or below the fitting (1.39 outliers were removed per model). For each fitting, we saved mean activity (Intercept), age effects, and edf. Next, we computed the derivative of each lifespan trajectory based on a numerical approximation with the *numDeriv* package. See clustering pipeline description in **Fig. 1**.

### Relationship between encoding activity, cognition, and GM variation

For each cluster, the first PCA component across ROIs was used as a participant’s episodic encoding measure. Cognition was assessed using: memory performance in the task, CVLT total learning score, and vocabulary and matrix reasoning raw scores. GM variation was derived from a linked ICA analysis (n = 70 components) using FLICA as described in (25) (http://fsl.fmrib.ox.ac.uk/fsl/fslwiki/FLICA) (**SI Methods and Results**). We then selected those components which showed a practical significance with age (r^2^ > .15). For each cluster, we fitted activity using GAM models with age and cognition/GM variation as smoothing terms and sex as a covariate. Note that these GAM analyses were run as implemented in the *mgcv* package as it allows for multiple smoothing terms and user-defined penalties to curve wiggliness (gamma = 2).

### Relationship between regional solution and topological organization

We assessed the topological relationship between cluster assignment and the first component of functional connectivity (3), flexibility (16), and macaque-human cortical expansion maps (15, 27). We ran a right-to-left registration for the evolutionary expansion map. Subcortical information was only available for the flexibility index. For the evolutionary expansion, we used ranked-values due to the exponential distribution. Values were averaged within ROIs. Permutation testing was used to establish the significance of topological relationships (n = 1000; random assignment of ROIs to clusters).

## Acknowledgments

This work was supported by the Department of Psychology, University of Oslo (to K.B.W., A.M.F.), the Norwegian Research Council (to K.B.W., A.M.F.) and the project has received funding from the European Research Council’s Starting Grant scheme under grant agreements 283634, 725025 (to A.M.F.) and 313440 (to K.B.W.).

