## Supplementary material for "The functional foundations of episodic memory remain stable throughout the lifespan": SI Methods

**Didac Vidal Piñeiro**

Department of Psychology, Pb. 1094 Blindern

Oslo, Norway, 0317

Tel: (+47) -22845089

**Keywords:**

Aging; development, encoding; episodic memory; fMRI.

### Supplementary Methods

#### Participants

The final sample included 540 individuals (females = 366, age = 39.1 [SD = 18.5], age range = 6-82). The participants were recruited from several projects coordinated by the Centre for Lifespan Changes in Brain and Cognition (LCBC, University of Oslo): The Norwegian Mother and Child Cohort Neurocognitive Study (1), Neurocognitive Development (2), Cognition and Plasticity Through the Lifespan (3), Constructive Memory (4), and Neurocognitive plasticity (5). All projects were approved by the Regional Ethical Committee of South Norway. All participants >12 years gave written informed consent, all participants <12 years gave oral informed consent and, for all participants <18 years, written informed consent was obtained from their legal guardians. All participants were screened through health and neuropsychological interviews. Initial exclusion criteria included neurologic or psychiatric disorders, chronic illness, premature birth, learning disabilities, left-handedness or, current use of medicines known to affect nervous system functioning. Participants were further excluded based on the following neuropsychological criteria: score <26 on the Mini-Mental State Examination (MMSE) (6), score of ≥21 in the Beck Depression Inventory (BDI) (7), score <85 on the Wechsler Abbreviated Scale of Intelligence (8), and a T-score of ≤30 on the California Verbal Learning Test II—Alternative Version (CVLT II) (9) immediate delay and long delay. Finally, participants' data were discarded due to technical errors, faulty acquisitions or a low number of trials in a condition of interest (< 6 trials; n = 14).

#### MRI acquisition

Imaging data were collected using a 20-channel head coil on a 3T MRI (Skyra, Siemens Medical Solutions, Ge) at Rikshospitalet (Oslo). Each encoding run consisted of 134 volumes with the following functional imaging parameters: 43 transversally oriented slices were measured using a BOLD-sensitive

T2*-weighted EPI sequence (TR = 2390 ms, TE = 30 ms, flip angle = $90^o$; voxel size = 3×3×3 mm; FOV = 224×224 mm; interleaved acquisition; generalized autocalibrating partially parallel acquisitions acceleration [GRAPPA] factor = 2). Three dummy volumes were collected at the start of each fMRI run to avoid T1 saturation effects in the analyzed data. Anatomical T1-weighted (T1w) magnetization-prepared rapid gradient echo (MPRAGE) images consisted of 176 sagittally oriented slices and were obtained using the following turbo field echo pulse sequence: TR = 2300 ms, TE = 2.98 ms, flip angle = $8^o$, voxel size = 1×1×1 mm, FOV = 256×256 mm. Additionally, a standard double-echo gradient-echo field map sequence was acquired for distortion correction of the echo-planar images. Visual stimuli were displayed in the scanner with an NNL 32-inch LCD monitor (1920×1080 pixels; NordicNeuroLab), positioned 176 cm from the mirror attached to the coil. Participants responded using the ResponseGrip system (NordicNeuroLab). Auditory stimuli were presented to the participants' headphones through the scanner intercom.

### Experimental design and behavioral analysis

This section describes in detail the experimental design illustrated in **Fig. 1a**. The stimulus material consisted of 300 black and white line drawings depicting everyday objects and items. The experiment consisted of an incidental encoding task and a surprise test, after ≈ 90 minutes, both inside the scanner. Only fMRI data during the encoding phase has been used in the present study. The encoding and the retrieval tasks consisted of two and four runs, respectively, that included 50 trials each. All runs started and ended with an 11 s baseline recording period in which a central fixation cross was present. An additional baseline period was also presented once in the middle of each run. In the encoding runs, a trial started with a prerecorded female voice asking through the participant's headphones, either "Can you eat it?" or "Can you lift it?" (in Norwegian). Each question was asked 25 times in each run in a pseudorandomized order. One second after the question onset, a picture of an item appeared on the screen (≈ 10 visual degrees in diameter) together with a response indicator that instructed the participant which button to press to respond “Yes” (the object can be eaten/lifted) or “No” (the object cannot be eaten/lifted). Button-response mapping was counterbalanced across participants. The subject had 2 s to produce a response before the object was replaced by a central fixation cross which remained on the screen throughout the intertrial interval (ITI), that lasted between 1-7 s (exponential distribution over four discrete ITIs; mean duration = 2.98 [SD 2.49] s). Despite the participants’ response-dependent nature of subsequent memory designs, the design efficiency – i.e. the distribution of ITIs in each encoding run - was tentatively optimized to ensure sufficient complexity in the recorded BOLD time series (http://surfer.nmr.mgh.harvard.edu/optseq/).

Participants were asked to perform a surprise memory test after ≈ 90 minutes of the last encoding trial. Test trials started with a recorded female voice asking the following (Q1): “Have you seen this item before”. Then, a picture of an item appeared on the screen, and the participant was instructed to indicate *Yes* (“I saw the item during the encoding phase”) or *No* (“I did not see the item during the encoding phase”) with a button press. In each run, 25 old and 25 new items were presented in a pseudorandomized order. Each object stayed on the screen for 2 s; if the participant responded that the item was new or did not respond, the trial ended. If the participant remembered seeing the item (pressed *Yes*), a new question followed (Q2): “Can you remember what you were supposed to do with the item?”. A *No* response ended the trial, whereas a *Yes* response, indicating that the participant also remembered the action associated with the item during the encoding, was followed by a final two-alternative forced-choice question (Q3): “Were you supposed to eat it or lift it?”. Here, the participant had to choose between the two actions “Eat” or “Lift” associated with the item encoding (“I imaged eating/lifting the item during the encoding phase”). The second question was included to discourage guessing behavior on the source memory question (Q3). The participants were verbally instructed minutes before both experimental tasks and did not go through any practice session before entering the scanner.

We computed additional behavioral measures in addition to source memory (Yes response to Q1 and Q2 and correct response to Q3); item memory (correct Yes response to Q1 and either a No response to Q2, or incorrect response to Q3) and miss (incorrect No response to Q1) – which are already specified in the main text (see *Experimental design and behavioral analysis* section). The additional measures corresponded to recognition hits (correct Yes response to Q1, regardless of response to Q2 and 3) and incorrect source judgments (incorrect eat/lift response to Q3). New items were classified either as correct rejections or false alarms. Memory performance in the task was assessed with a corrected source memory performance index (correct answers to Q3 - incorrect answers to Q3). This correction tentatively accounts for processes such as false memories, threshold criteria in Q2 or guessing behavior that affects the raw estimates of source memory performance (10, 11). fMRI conditions during encoding were modeled based on the behavioral response during the test phase.

### fMRI preprocessing

fMRI data were processed using the "fMRIPrep" preprocessing pipeline (12). Next, we detail the preprocessing steps based on the document generated by the fMRIPrep pipeline (v. 1.2.5). fMRIPrep is based on Nipype (v. 1.1.6) (13) while many of its internal operations use Nilearn 0.5.0 (14).

**Anatomical data preprocessing:** The T1w image was corrected for intensity non-uniformity (INU) using N4BiasFieldCorrection (15) (ANTs v. 2.2.0), and used as T1w-reference throughout the workflow. The T1w-reference was then skull-stripped using antsBrainExtraction.sh (ANTs v. 2.2.0), using OASIS as target template. Brain surfaces were reconstructed using recon-all (FreeSurfer v. 6.0.1) (16), and the brain mask estimated previously was refined with a custom variation of the method to reconcile ANTs-derived and FreeSurfer-derived segmentation of the cortical gray matter (GM) of Mindboggle (17). Spatial normalization to the ICBM 152 Nonlinear Asymmetrical template version 2009c (18) was performed through nonlinear registration with antsRegistration (ANTs v. 2.2.0) (19), using brain-extracted versions of both T1w volume and template. Brain tissue segmentation of cerebrospinal fluid (CSF), white-matter (WM) and GM was performed on the brain-extracted T1w using fast (FSL v. 5.0.9) (20).

**Functional data preprocessing:** For each BOLD run, the following preprocessing was performed. First, a reference volume and its skull-stripped version were generated using a custom methodology of fMRIPrep. A deformation field to correct for susceptibility distortions was estimated based on a field map that was co-registered to the BOLD reference, using a custom workflow of fMRIPrep derived from D. Greve's epidewarp.fsl script and further improvements of HCP Pipelines (21). Based on the estimated susceptibility distortion, an unwarped BOLD reference was calculated for a more accurate co-registration with the anatomical reference. The BOLD reference was then co-registered to the T1w reference using bbregister (FreeSurfer) which implements boundary-based registration (22). Co-registration was configured with six degrees of freedom. Head-motion parameters with respect to the BOLD reference (transformation matrices, and six corresponding rotation and translation parameters) are estimated before any spatiotemporal filtering using mcflirt (FSL v. 5.0.9) (23). BOLD runs were slice-time corrected using 3dTshift from AFNI v. 20160207 (24). The BOLD time-series (including slice-timing correction when applied) were resampled onto their original, native space by applying a single, composite transform to correct for head-motion and susceptibility distortions. These resampled BOLD time-series will be referred to as preprocessed BOLD in original space, or just preprocessed BOLD. Several confounding time-series were calculated based on the preprocessed BOLD: framewise displacement (FD) was calculated for each functional run, using Nipype's implementation (following the definitions by Power et al. (25). Additionally, a set of physiological regressors were extracted to allow for component-based noise correction (CompCor) (26). Principal components are estimated after high-pass filtering the preprocessed BOLD time-series (using a discrete cosine filter with 128s cut-off). A subcortical mask is obtained by heavily eroding the brain mask, which ensures it does not include cortical GM regions. Six anatomical CompCor (aCompCor) components are then calculated within the intersection of the aforementioned mask and the union of CSF and WM masks calculated in T1w space, after their projection to the native space of each functional run (using the inverse BOLD-to-T1w transformation). The head-motion estimates calculated in the correction step were also placed within the corresponding confounds file. All resamplings can be performed with a single interpolation step by composing all the pertinent transformations (i.e. head-motion transform matrices, susceptibility distortion correction when available, and co-registrations to anatomical and template spaces). Gridded (volumetric) resamplings were performed using antsApplyTransforms (ANTs), configured with Lanczos interpolation to minimize the smoothing effects of other kernels (27). Non-gridded (surface) resamplings were performed using mri_vol2surf (FreeSurfer).

### First-level fMRI analysis

First-level general linear models (GLM) were carried out with FSFAST (https://surfer.nmr.mgh.harvard.edu/fswiki/FsFast). For each participant and encoding run, we set up a first-level GLM consisting of the conditions of interest, with onsets and durations corresponding to the experimental trial period. GLMs were estimated both in the cortical surfaces and in the subcortical structures of interest in each subject's native space. Events were assigned to a given condition based on the participant's response during the subsequent memory test. The regressors were convolved with a double-gamma canonical hemodynamic response function (HRF). The conditions of interest were source and item memory conditions as defined in the behavioral analysis. Two additional regressors were included to soak up BOLD variance associated with miss memory trials and with trials with no response. Finally, for each participant, percent signal change during source and item memory encoding was estimated and contrasted to produce estimates of episodic memory encoding, which were then averaged over voxels/vertices within ROIs.

#### Linked ICA – Modes of GM variation analysis

To obtain modes of GM variation throughout the lifespan we used a linked Independent Component Analysis (ICA) as implemented in FLICA (http://fsl.fmrib.ox.ac.uk/fsl/fslwiki/FLICA) (28, 29) following the pipeline described by (30). The linked-ICA approach provides a data-driven decomposition of the images into spatial components characterizing the intersubject variability. Independent component approaches are able to model data into a set of – maximally independent; not orthogonal - interpretable features, some of them linked to biophysically plausible underlying mechanisms, which can additionally be linked to external variables such as age. Here, each spatial component represents a mode of variation of GM structure across n = 540 participants. We used three different modalities based on T1w-derived data: cortical thickness and area based on cortical surface reconstructions (16, 31, 32), and volume from a voxel-based morphometry (VBM) protocol (33, 34).

**Imaging processing.** Cortical thickness and cortical area maps were obtained through the FreeSurfer v.6.0 cortical reconstruction pipeline (http://surfer.nmr.mgh.harvard.edu/fswiki) (35–37). Briefly, the automatized processing pipeline feds on T1w images and includes removal of non-brain tissue, Talairach transformation, intensity correction, tissue and volumetric segmentation, cortical surface reconstruction and cortical parcellation. For each participant, the cortical surfaces were transformed into the *fsaverage5* template and smoothed by 12 mm (FWHM). GM volume maps were obtained through a FSL-VBM optimized protocol (33, 38, 39) (fsl.fmrib.ox.ac.uk/fsl/fslwiki/FSLVBM) (FSL v. 6.0.1). The volumes were initially masked by the full brain-segmented volume output from FreeSurfer obtained after nonuniformity correction. The images were then averaged and flipped along the x-axis to create a left-right symmetric, study-specific GM template. Next, the native GM images were non-linearly registered to the study-specific template and "modulated" to correct for local expansion/contraction due to the non-linear component of the spatial transformation. Finally, the modulated grey matter images were smoothed with an isotropic Gaussian kernel (σ = 3 mm; ≈7 FWHM).

**Linked ICA.** We ran the linked ICA decomposition – as implemented in FLICA (28, 29) - with 70 components as described in Douaud and colleagues (2014). For each independent component, we initially tested the relationship with age using generalized additive models (GAM; *mgcv* package; knots = 10; spline = "cr"; gamma = 2). After FDR-correction using the Benjamini–Yekutieli procedure (40) (pFDR), 20 components were significantly related to age (**Fig. S2**). Yet, only two reached *practical* significance; that is, explained 15% of the age variance. These two components, thereafter known as $IC_{GM}1$ and $IC_{GM}2$, were selected for further analyses. To assess the relative contribution of each modality in the independent components, we thresholded the weights at $Z > 4$.

### Vertexwise analysis across the entire sample

We carried a vertexwise analysis to display the subsequent memory contrast ($BOLD_{S>i}$: source vs. item memory encoding) cortical map for the entire sample across the cortical mantle (**Fig. 1c**). For each participant (n = 540), we transformed each fMRI regressor from the native to the *fsaverage6* template surface and obtained the $BOLD_{S>i}$ contrast by substracting the item to the source condition. Next, we concatenated the individual $BOLD_{S>i}$ maps and carried a one-sample t-test analysis using *mri_glmfit* (FreeSurfer). Statistical significance was considered at $p < .001$ FDR corrected bilaterally as implemented in *mri_fdr*.

### Supplementary Results

#### Linked ICA – Modes of GM variation analysis

While 20 components showed (pFDR < .05) significance with age (see **Fig. S2**), only two of them achieved a practical significance ($IC_{GM}1$ and $IC_{GM}2$; **Fig. 5**), which were consequently selected for further analysis. Both $IC_{GM}1$ and $IC_{GM}2$ replicate, to a great degree, the previous findings of (30). The $IC_{GM}1$ showed a global dominant mode of GM variation and exhibited a monotonic decrease across the lifespan. $IC_{GM}1$ was a multimodal component composed of cortical thickness (56%), volume (32%) and, cortical area (11%) information. Spatially, the $IC_{GM}1$ weighted strongly on large regions of the brain, both for the cortical thickness and the volume modalities. Contributions of cortical area information to $IC_{GM}1$ were limited to sensorimotor and occipital cortices (positive contributions in the gyri, negative in the sulci). $IC_{GM}2$ consisted also on a multimodal component determined to a great degree by cortical thickness information (61%) but also by cortical area and volume data (19% each). $IC_{GM}2$ weighted strongly on heteromodal frontoparietal networks in the cortical thickness and the volume modalities. In addition, medial temporal structures such as the hippocampus also contributed strongly to the component in the volume modality. The spatial pattern for the cortical area modality was relatively weak (thresholded at Z = 4) exhibiting positive effects in small frontoparietal regions and negative effects in lateral temporal and sensorimotor cortices. See representation in **Fig. S3**.

#### Vertexwise analysis across the entire sample

The BOLD correlates of subsequent episodic memory ($BOLD_{S>i}$) across the entire sample exhibited a canonical pattern of positive and negative subsequent memory effects (41). Widespread regions of the cortical mantle exhibited positive subsequent memory effects, particularly in the inferior and superior frontal gyrus, the medial temporal lobe, and in regions corresponding to the dorsal and ventral visual stream (including the fusiform, the isthmus cingulate and the superior parietal cortices).

The pattern was left-lateralized. Negative subsequent memory effects were mostly constrained to posteromedial and inferior parietal lobe regions, bilaterally. See **Fig. 1c** for a visual representation.

### Supplementary Figures

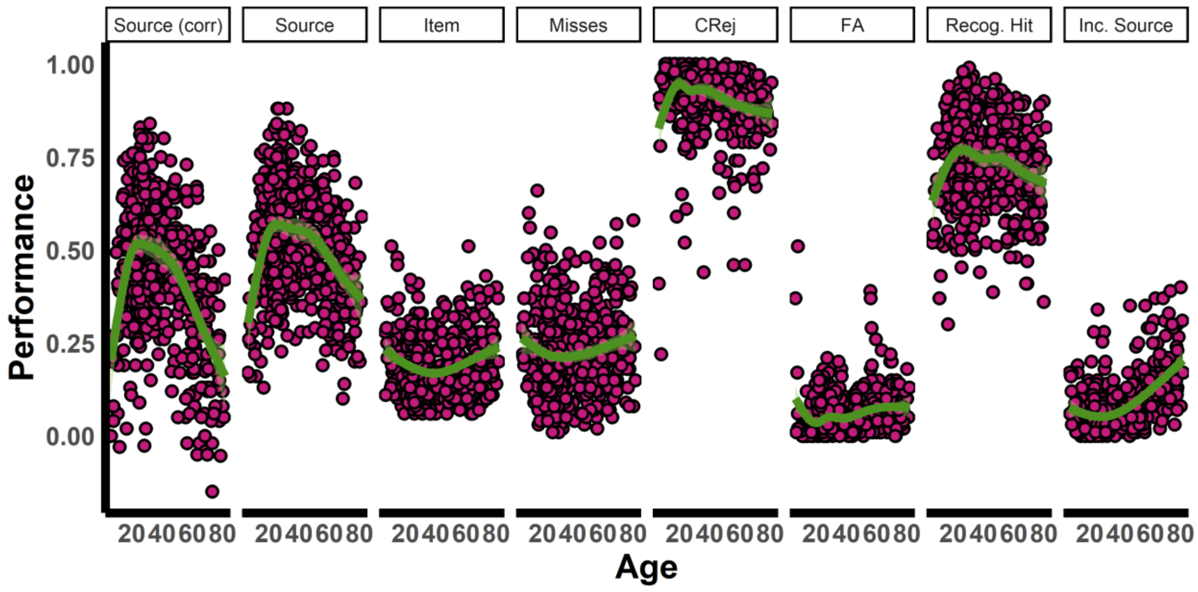


**Fig. S1.** ***Behavioral measures across the lifespan.*** *Smoothings are based on the GAM add-on implemented in ggplot2 (formula = y ~ s(x, bs = "cr", k = 8)). Values are proportional. See stats in* ***Table S1.*** *Note that the GAM stats* ***Table S1*** *are derived from mgcv. Source (corr) = Source hits (corrected); Source = Source hits; CRej = Correct rejections; FA = False alarms; Recog. Hit = Recognition hits; Inc. Source = Incorrect Source Judgments.*

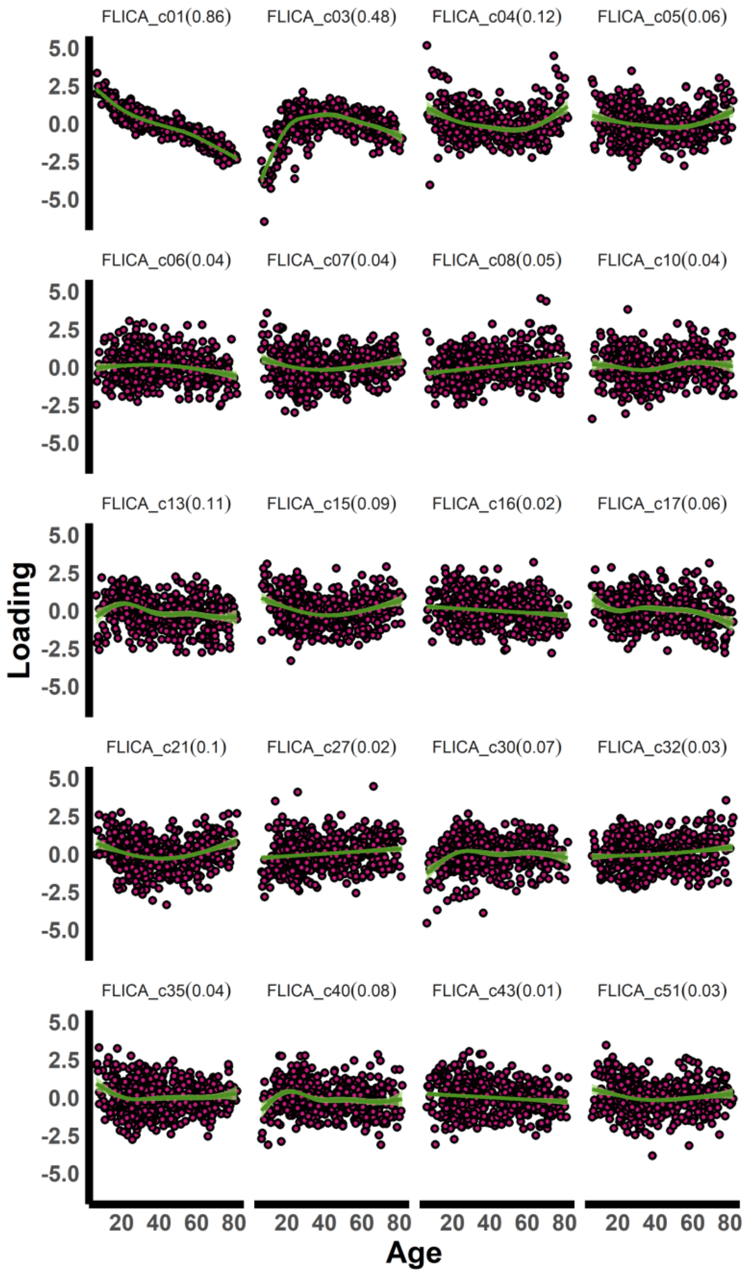


***Fig. S2. Linked-ICA components associated with age.*** *Relationship between Independent Components weights and Age. Only components with* pFDR < *.05 are shown. In parenthesis, age variance explained by the component (as assessed with GAM models). Only components «FLICA_01» and «FLICA_03» achieved practical significance and are hereafter referred to as* $IC_{GM}1$ *and* $IC_{GM}2$.

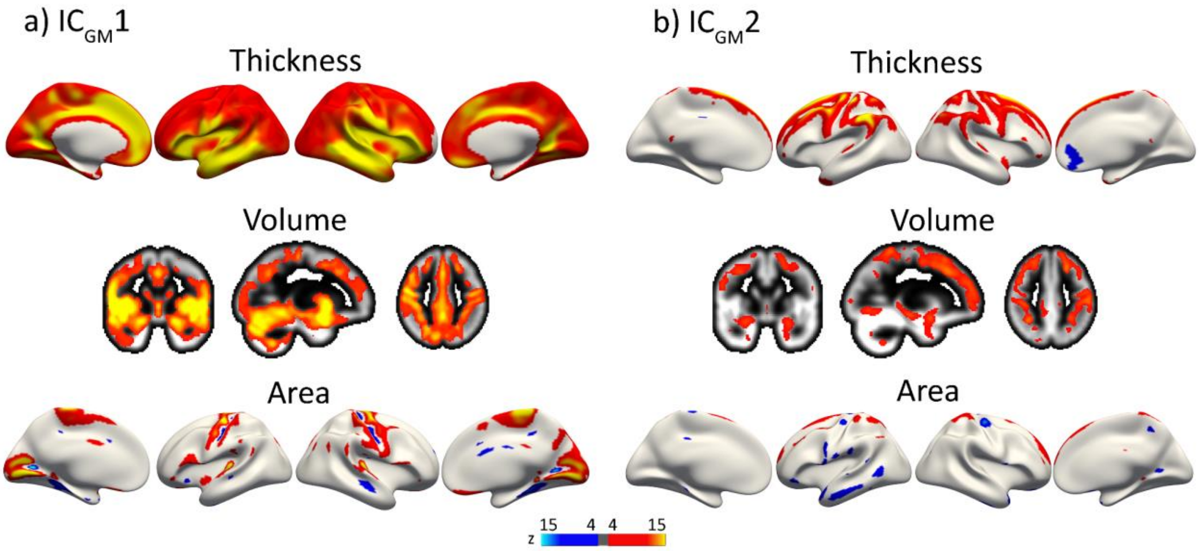


***Fig. S3. Spatial weight of independent GM components.*** *a, b) Modes of GM variation with practical ($r^2$ > .15) age significance. The current Figure expands the representation in* ***Fig. 5a,b*** *by including the weights for the cortical area modality and in the right hemisphere for cortical thickness.*

### Supplementary Tables

| | All | <15 | 15-20 | 20-30 | 30-40 | 40-50 | 50-60 | 60-70 | >70 | GAM stats |
|---|---|---|---|---|---|---|---|---|---|---|
| **N Participants** | 540 | 29 | 35 | 166 | 95 | 59 | 53 | 62 | 41 | --- |
| **Source hits (corrected)** | .44(.18) | .33(.20) | .47(.17) | .51(.15) | .51(.14) | .48(.14) | .38(.17) | .31(.18) | .22(.17) | 38.7(4.3; <.001)* |
| **Source hits** | .52(.15) | .41(.17) | .52(.15) | .56(.13) | .56(.13) | .56(.12) | .49(.13) | .45(.14) | .40(.12) | 20.4(4.2; <.001)* |
| **Item** | .19(.08) | .23(.12) | .19(.07) | .18(.07) | .17(.07) | .17(.08) | .20(.08) | .21(.09) | .22(.09) | 7(2.8; <.001)* |
| **Misses** | .23(.11) | .27(.13) | .23(.14) | .21(.11) | .22(.11) | .21(.10) | .24(.12) | .24(.12) | .25(.10) | 3.97(2.2; =.06) |
| **Correct rejections** | .92(.09) | .91(.17) | .94(.07) | .93(.07) | .93(.07) | .92(.07) | .89(.11) | .88(.09) | .87(.07) | 11(2.5; <.001)* |
| **False alarms** | .06(.06) | .06(.11) | .04(.04) | .05(.04) | .05(.04) | .05(.04) | .08(.09) | .08(.05) | .07(.05) | 14.7(1.2; <.001)* |
| **Recognition hits** | .74(.12) | .68(.13) | .74(.14) | .77(.11) | .76(.12) | .75(.11) | .73(.11) | .72(.12) | .70(.11) | 7.2(3.1; <.001)* |
| **Incorrect Source judgments** | .08(.07) | .08(.06) | .05(.04) | .06(.05) | .05(.04) | .08(.05) | .11(.07) | .14(.08) | .18(.09) | 64(3.2; <.001) |

***Table S1. Behavioral descriptives.*** *Mean (SD) of the behavioral measures derived from the fMRI memory task. Descriptives are shown for the entire sample and by age subgroups. Values are proportional. GAM stats indicate F(edf [estimated degrees of freedom]; pFDR). GAM was estimated with knots = 10, bs = "cr", method = "REML" and, gamma = 2. * denotes p < .05.*

| | CVLT learning | Matrices | Vocabulary | task perf. |
|---|---|---|---|---|
| **cl 0** | .62(1,1) | 2.56(3.5,.22) | .04(1,1) | .05(1,1) |
| **cl 1** | 1.78(1,.98) | 6.57(1,.1 | .65(1,1) | .81(1.4,1) |
| **cl 2** | .05(1.2,1) | .68(1,1) | 2.76(1,.59) | 4.18(1,.27) |
| **cl 3** | 5.21(1,.19 | 11.43(1.1,.02)* | 5.08(1.8,.06) | 3.82(2,.1) |
| **cl 4** | 8.55(1.3,.05) | 10.04(1.1,.04)* | 4.10(2.2,.08) | 7.78(2.2,.01)* |
| **cl 5** | 2.48(1,.66) | 7.02(1,.09 | 6.55(2.1,.01)* | 2.80(2.3,.22) |

***Table S2. Relationship between encoding clusters activity and cognition.*** *GAM stats assessing the relationship between cluster activity and cognition. GAM = F(edf, pFDR). GAM was estimated with knots = 10, bs = “cr”, method = “REML” and, gamma = 2. Age (as an additional smoothing term) and Sex were introduced as covariates. * denotes pFDR < .05. Matrices = Matrices Reasoning and Vocabulary = Vocabulary scores from Wechsler (1999). Task perf = task performance. CVLT learning = California Verbal Learning Test learning score* (9)*.*

| | $IC_{GM}1$ | $IC_{GM}2$ |
|---|---|---|
| **cl 0** | 2.91(1,.55) | 2.91(1,1) |
| **cl 1** | 0.39(1,1) | 0.39(1,1) |
| **cl 2** | 0.00(1,1) | 0.00(1,.55) |
| **cl 3** | 0.20(1,1) | 0.20(1,.36) |
| **cl 4** | 1.91(2,.55) | 1.91(1,.01)* |
| **cl 5** | 5.70(2.5,.01)* | 5.70(1,<.001)* |

***Table S3. Relationship between encoding clusters activity and GM variation.*** *GAM stats assessing the relationship between cluster activity and GM variation. GAM = F(edf, FDRp). GAM was estimated with knots = 10, bs = "cr", method = "REML" and, gamma = 2. Age (as an additional smoothing term) and Sex were introduced as covariates. * denotes pFDR < .05.*

| | Flexibility | Conn. Gradient | Expansion |
|---|---|---|---|
| **cl 0** | 1.58(1.00); 1 | -2.36(3.22); 1 | .61(.15); 1 |
| **cl 1** | 1.66(1.08); 1 | -0.12(3.93);1 | .61(.18); 1 |
| **cl 2** | 1.27(0.93); 1 | -1.46(3.66);1 | .57(.16); 1 |
| **cl 3** | 2.16(1.54); .06 | 2.4(3.40); >.001* | .66(.15); .28 |
| **cl 4** | 2.86(1.51); .002* | 3.72(1.77); >.001* | .71(.13); .04* |
| **cl 5** | 1.63(0.80); 1 | -0.48(3.11); 1 | .47(.21); 1 |

***Table S4. Topological relationship between encoding clusters and functional and evolutionary hierarchy.*** *Mean (SD); FDRp values for each functional/evolutionary hierarchy, grouped by cluster. Observations correspond to each of the |N| = 416 ROIs. Conn. Gradient = principal gradient of functional connectivity. * denotes pFDR < .05 using permutation testing.*
